# X chromosome factor Kdm6a enhances memory independent of its demethylase function in the aging XY male brain

**DOI:** 10.1101/2022.07.18.500498

**Authors:** Cayce K. Shaw, Samira Abdulai-Saiku, Francesca Marino, Dan Wang, Emily J. Davis, Barbara Panning, Dena B. Dubal

**Affiliations:** Department of Neurology and Weill Institute for Neurosciences, University of California, San Francisco, CA 94158; Rehabilitation Sciences Graduate Program, University of California, San Francisco, CA 94158; Neuroscience Graduate Program, University of California, San Francisco, CA 94158; Department of Biochemistry and Biophysics, San Francisco, CA 94158

**Author notes:** Correspondence (D.B.D.), 675 Nelson Rising Lane, Rm 212B, San Francisco, CA 94110.

## Abstract

Males exhibit shorter lifespan and more cognitive deficits in aging human populations. In mammals, the X chromosome is enriched for neural genes and is a major source of biologic sex difference, in part, because males show decreased expression of select X factors. While each sex (XX and XY) harbors one active X due to X chromosome inactivation in females, some genes, such as *Kdm6a*, transcriptionally escape silencing in females – resulting in lower levels in males. Kdm6a is a known histone demethylase (H3K27me2/3) with multiple functional domains that is linked with synaptic plasticity and cognition. Whether elevating *Kdm6a* could benefit the aging male brain and whether this requires its demethylase function remains unknown. We used lentiviral-mediated overexpression of the X factor in the hippocampus of aging male mice and tested their cognition and behavior in the Morris water maze. We found that acutely increasing *Kdm6a* – in a form without demethylase function – selectively improved learning and memory, without altering total activity or anxiety-like measures, in the aging XY brain. Further understanding the demethylase-independent downstream mechanisms of Kdm6a may lead to novel therapies for treating age-induced cognitive deficits in both sexes.

---

Males exhibit shorter lifespan [1, 2] and more cognitive deficits [3-5] in aging human populations. Sex chromosomes, and specifically the X chromosome, are a major source of sex difference, such as in lifespan [6]. X-derived, sex difference can result from decreased expression of select X-linked factors in males. The X chromosome is enriched for neural genes and, in females, associates with less cognitive decline in aging [7]. While both males (XY) and females (XX) harbor one active X due to X chromosome inactivation (XCI) in females [8], some genes escape transcriptional silencing – resulting in comparatively lower levels in males [9]. Since cognitive decline is a major biomedical challenge, understanding whether and how specific X factors promote brain health could open avenues for novel treatments for both sexes.

Kdm6a, or lysine demethylase 6a, is an X factor that influences synaptic plasticity [10] and cognition [10-12] and, due to XCI, exhibits lower levels in XY human and mouse brains [12-14]. Kdm6a contains several functional domains and is primarily known for its nuclear histone demethylase (H3K27me2/3) activity [15-17], though its location in hippocampal neurons is largely cytoplasmic [12]. *Kdm6a* is highly enriched in the brains of humans and mice [12, 18]. In humans, genetic variation leading to higher *KDM6A* levels in the brain associates with less cognitive decline in both sexes [12]. In XY mice, *Kdm6a* elevation in the hippocampus confers resilience against Alzheimer’s disease-related toxicity [12].

Whether elevating *Kdm6a* could benefit the aging male brain, and whether this requires its demethylase function remains unknown. Here, we show that acutely increasing *Kdm6a* – in a form without demethylase function [19, 20] – selectively improved memory, without altering total activity or anxiety-like measures, in the aging XY brain.

## MATERIALS AND METHODS

### Experimental Animals

Mice for *in vivo* studies were on a congenic C57BL/6J background and kept on a 12-hr light/dark cycle with ad libitum access to food and water. The standard housing group was five mice per cage except for single housing during water maze studies. Cognitive and behavioral studies were carried out during the light cycle. Studies were approved by the Institutional Animal Care and Use Committee of the University of California, San Francisco, and conducted in compliance with National Institutes of Health guidelines. All cognitive, behavioral, and molecular experiments were conducted in a blinded manner on age-matched littermate offspring.

### Lentivirus production and stereotaxic injection

The *Kdm6a* Enzyme-Dead (Kdm6a DeM-dead) plasmid was purified and validated by Addgene (#40619) in which alanine (A) substitutions were made at histidine (H) 1146 [19] and glutamic acid (E) 1148 [19] of a sequence encoding protein, Kdm6a (NCBI Reference Sequence: NM_009483.2; 4275 bp). This sequence was inserted between AscI and BmtI restriction sites of the pSicoR lentiviral backbone (pSicoR-EF1a-Blast-T2A-EGFP) obtained from the UCSF ViraCore (catalog number: MP394). Kdm6a cDNA is ∼4kB in size. Active lentiviral particles were produced as previously described [12].

Male mice, 17-to 18-month-old, were anesthetized using isofluorane at 2-3% and placed in a stereotaxic frame. Lentiviral vectors of Kdm6a DeM-dead or control virus (5 uL per hemisphere, MOI = 2) were stereotactically injected bilaterally into the dentate gyrus of the hippocampus using the coordinates AP=-2.1, ML=±1.7 and DV=1.9. All behavioral assays were conducted 3-5 weeks after lentiviral injections.

### General cell culture

Primary cortical cell cultures were isolated from postnatal days 0-2 congenic C57BL/6J XY mouse pups as described [12]. Cells were plated at 1 million cells/mL in 24-well plates for subsequent maturation and treatment in neurobasal media with B-27 supplement (NBA/B27). At day *in vitro* (DIV) 4, cultures were transduced with Kdm6a DeM-dead lentivirus at MOI=2. Cells were harvested and RNA isolated at times indicated.

### Quantitative PCR

Quantitative PCR was performed on RNA isolated from *in vitro* neurons and *in vivo* dentate gyrus. To verify Kdm6a DeM-dead lentiviral transfection, the dentate gyrus was dissected from the whole hippocampus under a stereomicroscope (Zeiss, Stemi 2000-C) in RNase-free conditions. The RNA was then isolated using Sigma Aldrich RNAzol RT (Cat. R4533). Real-time PCR of *18S* and *Kdm6a* was performed. Primers for *Kdm6a* exons 16-17 5’- ATAACCGCACAAACCTGACC and 5’- ACCTGCCAAATGTGAACTCG were used to measure *Kdm6a* mRNA expression.

### Elevated plus-maze

Testing was carried out as described [12, 21, 22]. Briefly, mice were habituated to the testing room for 1 hour prior to testing. Dim light was maintained in the testing room for both habituation and testing. Mice were placed in the center of an elevated plus-maze facing an open arm and allowed to explore for 10 minutes. Total time spent in open and closed arms was recorded using Kinder Scientific Elevated Plus-Maze and MotorMonitor™ system.

### Open field

Testing was carried out as described [12, 21, 22]. Briefly, mice were acclimated to the room for 30 minutes and allowed to explore the open field for 5 minutes. Total activity in the open field (clear plastic chamber, 41 × 30 cm) was detected by beam breaks and measured with an automated Flex-Field/Open Field Photobeam Activity System (San Diego Instruments).

### Morris water-maze

Testing was carried out as described [12, 21-23]. Briefly, the water maze pool (diameter, 122 cm) contained white, opaque water with a square, 14 cm^2^ platform submerged 2 cm below the surface. During hidden platform training, the platform location remained constant, and the drop location varied between trials. Mice received two training sessions, consisting of two trials each, daily for seven days. The maximum time allowed per trial was 60 seconds. For the probe trial, the platform was removed, and the mice were allowed 60 seconds to swim. Following probe testing, mice were tested for their ability to find the platform when marked with a visible cue (15 cm pole placed on the platform).

### Statistical analyses

GraphPad Prism (version 7.0) was used for *t* tests and visualization of data. R (nmle package) was used for analyses of variance (ANOVAs) and post hoc tests. Differences between two means were assessed by two-tailed *t* tests unless otherwise indicated. Differences among multiple means were assessed by two-way ANOVA. A mixed-model ANOVA was used for analyses of Morris water-maze data and included effects of repeated measures. Only significant *P* values were stated for two-way ANOVA results. Multiple comparisons of post-hoc *t* tests were corrected for with the Bonferroni-Holm (stepwise Bonferroni) procedure to control for a family-wise error rate of α = 0.05. Exclusion criteria (greater than 2 SDs above or below the mean) were defined a priori to ensure unbiased exclusion of outliers. Error bars represent ± SEM. Null hypotheses were rejected at or below a *P* value of 0.05.

## RESULTS

### *Kdm6a* is decreased in the hippocampus of old XY mice, and its demethylase-dead form (Kdm6a DeM-Dead) was overexpressed in aged XY brains

We first assessed whether sex bias in *Kdm6a* mRNA levels extends to the aging hippocampus. Indeed, *Kdm6a* levels were significantly lower in the aging male hippocampus (Fig. 1A). To test if elevating *Kdm6a* could benefit the aging XY brain in a histone demethylase (H3K27me2/3) independent manner, we utilized a *Kdm6a* lentivirus construct containing point mutations at the catalytic jumonji-C domain (H1146A/E1148A), rendering dead its demethylase function as previously demonstrated [19] (Fig. 1B). The Kdm6a demethylase-dead (Kdm6a DeM-dead) construct increased *Kdm6a* mRNA expression levels via lentiviral-mediated transfection in XY primary neurons compared to the control (Fig. 1C-D).

**Figure 1.**
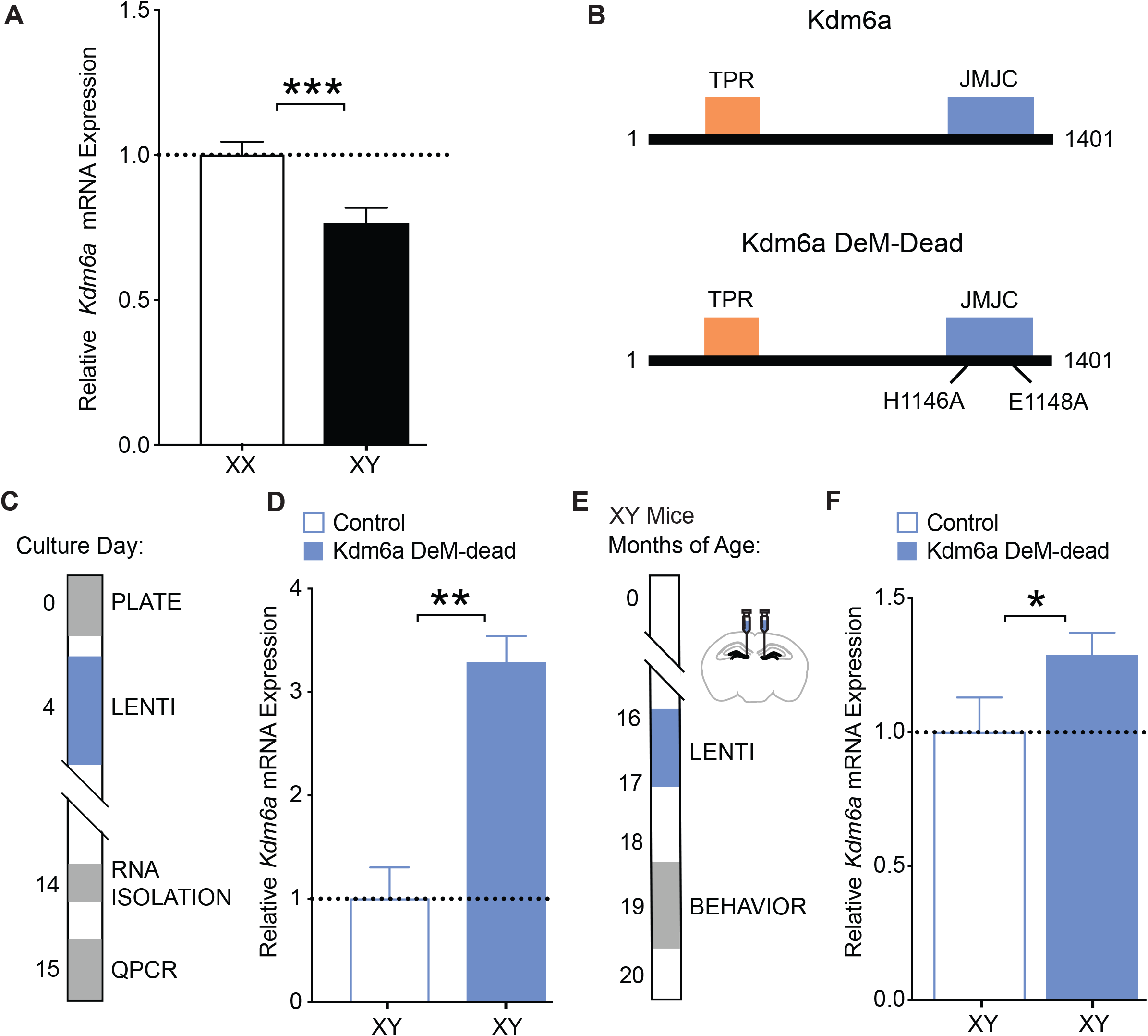
*Kdm6a* is decreased in the hippocampus of aged XY mice, and its demethylase-dead form (Kdm6a Dem-Dead) was overexpressed in aged XY brains. (A) Hippocampal *Kdm6a* mRNA expression in aged XX and XY mice (age 20-35 months; n=10-13 mice per group, shown relative to XX mice). (B) Construct maps showing the mutations rendering dead the demethylase activity of Kdm6a (Kdm6a DeM-Dead) along with the tetrapeptide repeat (TPR) domain. (C) Experimental strategy of lentivirus-mediated overexpression of Kdm6a DeM-Dead in XY mouse primary cortical neurons. (D) *Kdm6a* mRNA levels in primary XY neurons transfected with lentivirus expressing control or Kdm6a DeM-dead, shown relative to control (n=3 wells per experimental group from 10 XY pups). (E) Experimental strategy of lentiviral injection followed by testing in behavioral tasks. (F) *Kdm6a* mRNA expression following lentiviral transfection of Kdm6a DeM-Dead measured in the dentate gyrus of the hippocampus, relative to XY controls (n=3 mice per group) *P < 0.05 (two-tailed *t* tests in A, D; one-tailed in F). Data are presented as means ± SEM.

We next increased Kdm6a DeM-dead expression in the hippocampus of aging XY mice. We injected lentivirus with (Kdm6a DeM-dead) or without (control) the *Kdm6a* construct bilaterally into the dentate gyrus, a region that affects spatial learning and memory, and analyzed mice behaviorally 1 month later (Fig. 1E). Lentiviral-mediated overexpression of Kdm6a DeM-dead increased *Kdm6a* mRNA expression in the dentate gyrus (Fig. 1F).

### *Kdm6a* enhances learning and memory, independent of its demethylase function, in aging XY male mice

We tested XY mice in cognitive and behavioral tasks (Fig. 2A) to determine if elevating *Kdm6a* mRNA levels, in its demethylase dead form, could improve learning and memory. Increasing Kdm6a DeM-dead expression in XY aged mice did not alter time spent in open arms (Fig. 2B) in the elevated plus-maze or movements in the open field task (Fig. 2C), indicating no changes in anxiety-like behavior or total activity. In contrast, increasing Kdm6a DeM-dead expression in XY aged mice improved learning in the Morris water maze, measured by shorter distance traveled to the hidden platform (Fig. 2D). Additionally, in probe trials, increasing XY Kdm6a DeM-dead robustly improved spatial memory, compared to XY control mice (Fig. 2E). Swim speeds and distance to find a visible cue, experimental controls, did not differ between the groups (data not shown). Thus, overexpression of *Kdm6a* in its demethylase dead form selectively improved learning and memory, without altering total activity or anxiety-like measures, in the aging XY brain.

**Figure 2.**
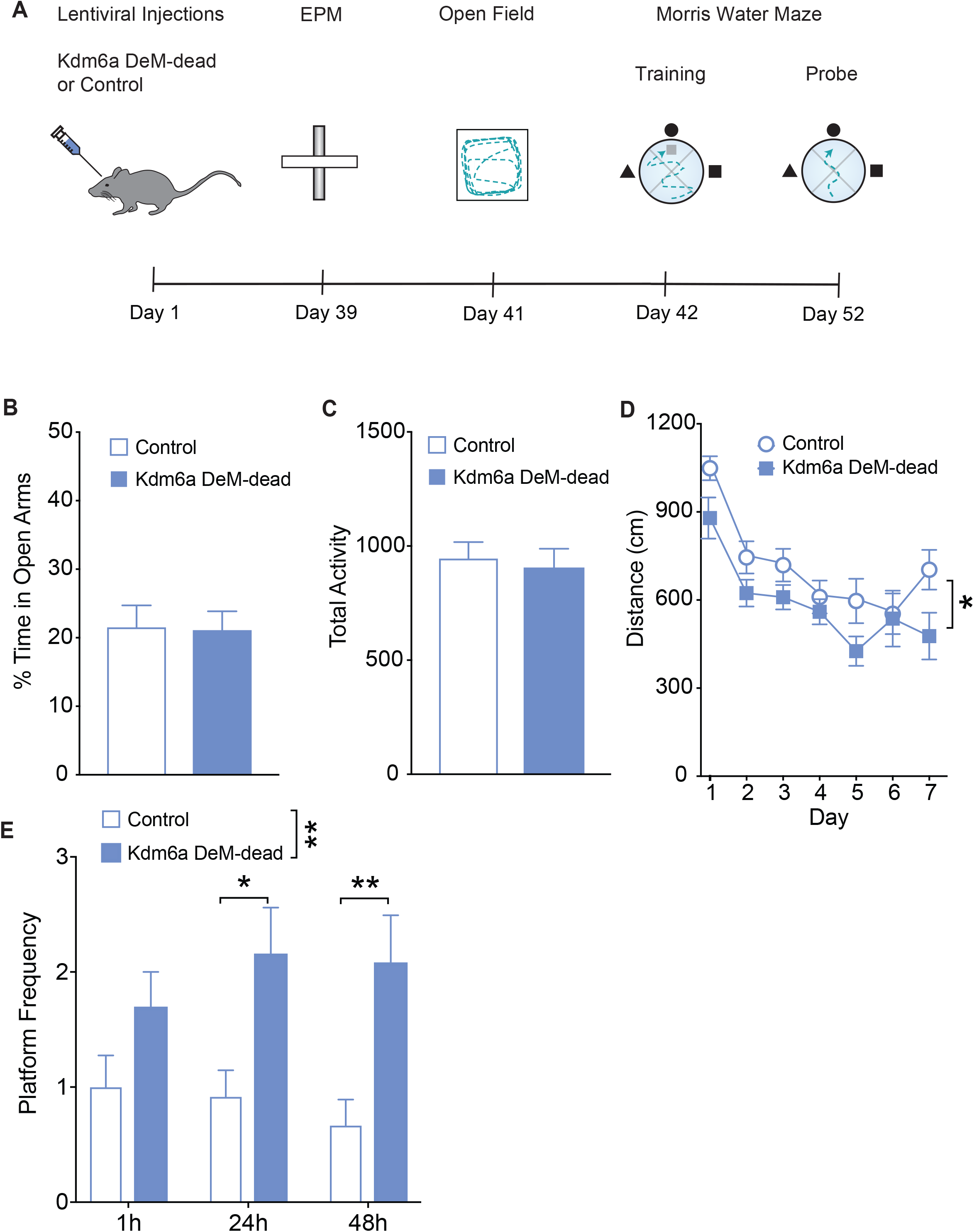
Kdm6a demethylase-dead overexpression in hippocampus improves cognition in XY aged mice. (A) Diagram of the experimental strategy for cognitive testing in the elevated plus-maze, open field testing, and the Morris water maze in aged XY mice (age 17-20 months, n=13-14 mice per experimental group). (B) Percentage of time spent in the open arms of the elevated plus-maze during 10 min exploration period. (C) Total number of movements during exploration of the open field for 5 min. (D) Spatial learning curves (platform hidden) of aged mice, control or Kdm6a DeM-dead, in the Morris water maze. Overexpressing Kdm6a DeM-dead mRNA enhanced learning. Two-way ANOVA: treatment *P < 0.05. (E) Probe trial results 1 hour, 24 hours, and 48 hours after completion of hidden platform learning, indicating spatial memory, showed that Kdm6a DeM-dead overexpressing mice had attenuated spatial deficits measured by increased frequency of entries into the target zone, compared to control mice. Two-way ANOVA: treatment *P < 0.05; **P < 0.01 (Bonferroni-Holm). Data are presented as means + SEM.

## DISCUSSION

Our studies in mice collectively reveal that *Kdm6a* is decreased in the aging XY hippocampus and overexpressing it in the dentate gyrus – in a form without demethylase function – selectively improved learning and memory in aging XY mice. These data support the hypothesis that elevating Kdm6a enhances the brain in aging.

An acute and modest increase of *Kdm6a*, without its demethylase activity, in the dentate gyrus of aged XY mice improved learning and memory. Whether the extent of improvement could match wildtype increases of *Kdm6a* is not known since this was not tested in parallel. It is noteworthy that a small increase, similar to or less than levels found in females, was sufficient to improve learning and memory in males. This suggests that lower levels of *Kdm6a* in males, compared to females, may confer vulnerability to cognitive aging. It remains unknown whether even higher levels could achieve further protection – and whether increasing *Kdm6a* in females, beyond its endogenously higher levels, could also improve cognition in aging.

The small *Kdm6a* increase in males occurred during the old life stage and acutely countered cognitive deficits present at this age, suggesting that Kdm6a manipulations, even late in life, could be beneficial. Consistent with findings that manipulating cells within functional hubs can affect larger networks [24], increasing *Kdm6a* levels in a small subregion of the XY hippocampus improved cognition. It will be important to investigate pharmacologic or lifestyle pathways to increase *Kdm6a* in the brains of both sexes.

In our studies, the lack of demethylase function in Kdm6a-mediated cognitive improvement in males indicates that other yet-identified functions of Kdm6a improve learning and memory. For example, Kdm6a can act as a scaffold for a larger transcription factor complex based on its tetratricopeptide repeat (TPR) domain [25, 26]. Kdm6a can also indirectly regulate levels of H3K27 methylation by controlling enhancer activity of other demethylase enzymes [26]. Future studies will identify specific mechanisms of Kdm6a demethylase-independent functions in cognition.

Strategies for targeting demethylase-independent mechanisms of Kdm6a in the brain could potentially lead to novel therapies for treating cognitive deficits in aging males, females, or both.

## ACKNOWLEDGEMENTS

We thank Chen Chen for assistance with mouse colony maintenance.

## REFERENCES

1. United Nations, D.o.E.a.S.A., Population Division, World Population Ageing 2019. 2020.

2. Zarulli, V., et al., Women live longer than men even during severe famines and epidemics. Proceedings of the National Academy of Sciences, 2018. 115(4): p. E832–E840.

3. Casaletto, K.B., et al., Cognitive aging is not created equally: differentiating unique cognitive phenotypes in “normal” adults. Neurobiol Aging, 2019. 77: p. 13–19.

4. Jack, C.R., Jr., et al., Age, Sex, and APOE ε4 Effects on Memory, Brain Structure, and β-Amyloid Across the Adult Life Span. JAMA Neurol, 2015. 72(5): p. 511–9.

5. Dubal, D.B., Sex difference in Alzheimer’s disease: An updated, balanced and emerging perspective on differing vulnerabilities. Handb Clin Neurol, 2020. 175: p. 261–273.

6. Davis, E.J., I. Lobach, and D.B. Dubal, Female XX sex chromosomes increase survival and extend lifespan in aging mice. Aging Cell, 2019. 18(1): p. e12871.

7. Davis, E.J., et al., Sex-Specific Association of the X Chromosome With Cognitive Change and Tau Pathology in Aging and Alzheimer Disease. JAMA Neurol, 2021. 78(10): p. 1249–1254.

8. Lyon, M.F., Gene action in the X-chromosome of the mouse (Mus musculus L.). Nature, 1961. 190: p. 372–3.

9. Berletch, J.B., F. Yang, and C.M. Disteche, Escape from X inactivation in mice and humans. Genome Biol, 2010. 11(6): p. 213.

10. Tang, G.B., et al., The Histone H3K27 Demethylase UTX Regulates Synaptic Plasticity and Cognitive Behaviors in Mice. Front Mol Neurosci, 2017. 10: p. 267.

11. Faundes, V., et al., Clinical delineation, sex differences, and genotype-phenotype correlation in pathogenic KDM6A variants causing X-linked Kabuki syndrome type 2. Genet Med, 2021. 23(7): p. 1202–1210.

12. Davis, E.J., et al., A second X chromosome contributes to resilience in a mouse model of Alzheimer’s disease. Sci Transl Med, 2020. 12(558).

13. Greenfield, A., et al., The UTX gene escapes X inactivation in mice and humans. Hum Mol Genet, 1998. 7(4): p. 737–42.

14. Berletch, J.B., et al., Identification of genes escaping X inactivation by allelic expression analysis in a novel hybrid mouse model. Data Brief, 2015. 5: p. 761–9.

15. Gažová, I., A. Lengeling, and K.M. Summers, Lysine demethylases KDM6A and UTY: The X and Y of histone demethylation. Mol Genet Metab, 2019. 127(1): p. 31–44.

16. Sengoku, T. and S. Yokoyama, Structural basis for histone H3 Lys 27 demethylation by UTX/KDM6A. Genes Dev, 2011. 25(21): p. 2266–77.

17. Agger, K., et al., UTX and JMJD3 are histone H3K27 demethylases involved in HOX gene regulation and development. Nature, 2007. 449(7163): p. 731–4.

18. Xu, J., et al., Sex-specific differences in expression of histone demethylases Utx and Uty in mouse brain and neurons. J Neurosci, 2008. 28(17): p. 4521–7.

19. Wang, C., et al., UTX regulates mesoderm differentiation of embryonic stem cells independent of H3K27 demethylase activity. Proc Natl Acad Sci U S A, 2012. 109(38): p. 15324–9.

20. Yoo, K.H., et al., Histone Demethylase KDM6A Controls the Mammary Luminal Lineage through Enzyme-Independent Mechanisms. Mol Cell Biol, 2016. 36(16): p. 2108–20.

21. Dubal, D.B., et al., Life extension factor klotho enhances cognition. Cell Rep, 2014. 7(4): p. 1065–76.

22. Dubal, D.B., et al., Life extension factor klotho prevents mortality and enhances cognition in hAPP transgenic mice. J Neurosci, 2015. 35(6): p. 2358–71.

23. Gupta, S., et al., KL1 Domain of Longevity Factor Klotho Mimics the Metabolome of Cognitive Stimulation and Enhances Cognition in Young and Aging Mice. J Neurosci, 2022. 42(19): p. 4016–4025.

24. Bonifazi, P., et al., GABAergic hub neurons orchestrate synchrony in developing hippocampal networks. Science, 2009. 326(5958): p. 1419–24.

25. Van der Meulen, J., F. Speleman, and P. Van Vlierberghe, The H3K27me3 demethylase UTX in normal development and disease. Epigenetics, 2014. 9(5): p. 658–668.

26. Wang, S.P., et al., A UTX-MLL4-p300 Transcriptional Regulatory Network Coordinately Shapes Active Enhancer Landscapes for Eliciting Transcription. Mol Cell, 2017. 67(2): p. 308-321.e6.

